# Carcinogenesis is associated with sex-biased tissue environments and imbalances between estrogen receptor alpha and thyroid hormone receptors

**DOI:** 10.1101/2023.05.20.541565

**Authors:** Feng Zhang, Minghua Deng

**Affiliations:** Center for Quantitative Biology, Academy for Advanced Interdisciplinary Studies, Peking University, Beijing, China; School of Mathematical Sciences, Peking University, Beijing, China; Center for Statistical Science, Peking University, Beijing, China

## Abstract

There is extensive evidence of sex differences in the susceptibility and prognosis of non-reproductive cancers. In addition to external factors, biological sex bias (e.g., sex chromosomes, hormones, and immune function) is suspected to function as a selective pressure that influences the evolutionary process of carcinogenesis. However, there remains a lack of clarity concerning the extent of the effect of sex bias on carcinogenesis, as well as the underlying mechanism. In this study, we show that tissue sex bias, correlated to gonadotropin-releasing hormone, varies among tissues and is associated with two distinct age-specific patterns of cancer incidence: parallel and nonparallel. Additionally, we reveal that imbalances in estrogen receptor alpha and thyroid hormone receptors are associated with levels of hypoxia-inducible factors, which have three phases (hypoxia, hyperoxia, and “chaotic-oxia”) that exist in most cancers and are linked to specific cancer subtypes, including cancers with microsatellites, the CpG island methylator phenotype, or hypermethylation. Our results suggest that sex-biased tissue environments and hormonal imbalances may influence the incidence pattern and direction of carcinogenesis, emphasizing the importance of maintaining hormonal homeostasis for cancer prevention and providing insights toward improving therapies for cancer types with hormonal imbalances.

**Plain English summary:** Sex bias has been long observed in cancer susceptibility and prognosis. Hormone difference between the two sexes was believed to play a role. However, the mechanism is still largely unknown. Our study has shown that the activity of one hormone secreted from pituitary, namely gonadotropin-releasing hormone, are correlated to sex-biased tissue environment, which was speculated to influence incidence patterns of carcinogenesis. Furthermore, hormonal imbalance, particularly imbalance in receptors of estrogen and thyroid hormone, which is associated with specific cancer subtypes, was speculated to influence the direction of carcinogenesis.

**Highlights:**

- There are two distinct patterns of cancer age-specific incidence curve between the two sexes: parallel and nonparallel.
- The parallel and nonparallel patterns are associated with sex bias in tissue environments which is correlated with sex-biased activity of gonadotropin-releasing hormone.
- Three sex-biased mutated genes: *PTEN*, *PGM5* and *LARP4B*, have tissue-specific hotspot mutations which are associated with microsatellite instability (MSI), and the proportion of MSI exhibit sex bias in stomach cancer.
- Three hypoxia induce factors (HIFs) phases: hypoxia, hyperoxia, and “chaotic-oxia”, exist in most cancers and are linked to specific cancer subtypes, including cancers with MSI, the CpG island methylator phenotype, and hypermethylation.
- The phases of HIFs are associated with imbalances between estrogen receptor alpha and thyroid hormone receptors.

## Introduction

The age-specific incidence curve of a particular cancer can represent the evolutionary dynamics of carcinogenesis. For carcinomas (i.e., most solid cancers), age-specific incidence curves are approximately linear on a log-log scale (1). Armitage and Doll proposed a multistage model explaining the power law relationship, based on the assumption that carcinogenesis is a multistage process involving the accumulation of driver mutations (2). This model has been modified by other studies to incorporate new information (3–8). Carcinogenesis is influenced by selective pressure from the microenvironment, through a process in which driver mutations (genetic or epigenetic) are selected and accumulated. Generally, different cancer types have distinct mutational spectra (9, 10), indicating that variations in tissue environment and cell type can influence the evolutionary dynamics of carcinogenesis.

For nearly all non-reproductive cancers, women and men have different susceptibilities and prognoses. However, behavior or mutagen exposure can only partly explain this phenomenon (11). Biological sex bias, involving sex chromosomes and hormones, also plays a role in many instances (12–15). Recent studies have shown that female and male patients preferentially exhibit mutations of certain genes and gain specific transcriptional or epigenetic signatures (16–18); because they display stronger immunity, female patients preferentially accumulate more mutant peptides that are not recognized by T cell receptors (19). These studies provide evidence to support the hypothesis that sex bias is a selective pressure with the potential to influence carcinogenesis. However, the mechanism by which sex bias influences carcinogenesis remains largely unknown.

## Results

### Parallel and nonparallel patterns of age-specific cancer incidence in women and men

Based on the hypothesis that sex bias influences carcinogenesis which can be represented by the age-specific incidence curve of a particular cancer, we used the CI5-X database (provided by the International Agency for Research on Cancer) to compare the incidence curves of major solid cancer types between women and men in selected regions or ethnicities (see methods). Fitted age-specific incidence curves for men and women showed two general patterns on a log-log scale: parallel and nonparallel (Fig. 1A). Parallel patterns were observed when colon cancer incidences were compared between inherited and sporadic types (7), and when lung cancer incidences were compared between smokers and non-smokers (20). These parallel patterns suggest that individuals with a susceptible genetic background or greater exposure to mutagens have a higher baseline risk of cancer. To our knowledge, nonparallel patterns have not been reported.

**Fig. 1:**
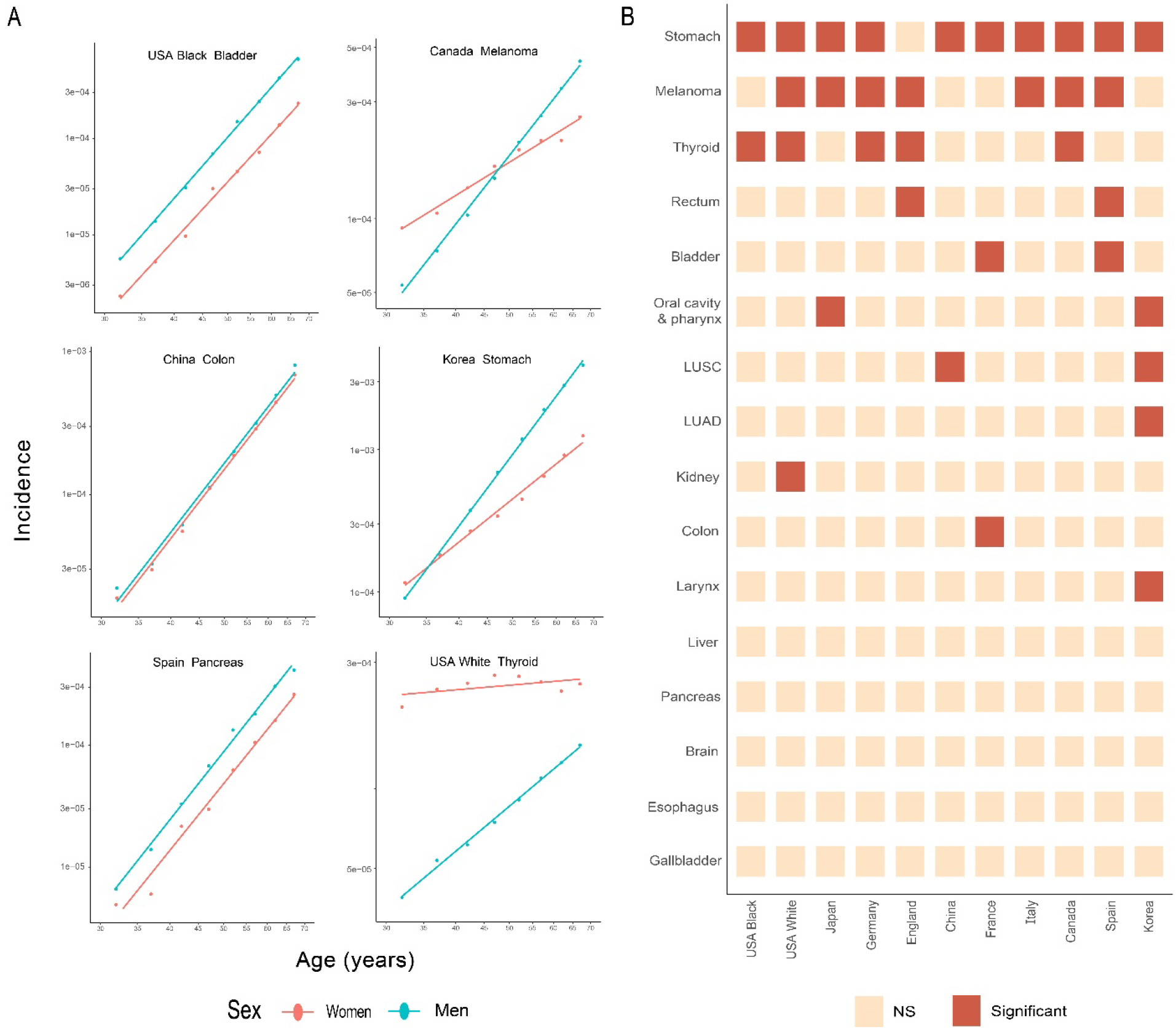
Cancer incidence curves and prevalence of nonparallel patterns. (A) Examples of parallel and nonparallel patterns. Both axes are on a logarithmic scale. Parallel patterns are shown on the left; nonparallel patterns are shown on the right. Incidences and fitted curves for women are shown in red; incidences and fitted curves for men are shown in blue. (B) Prevalence of significant nonparallel patterns in major cancer types according to region or ethnicity. Significant nonparallel patterns are shown in red; non-significant (NS) nonparallel patterns are shown in light yellow.

To determine the prevalence of nonparallel patterns in major solid cancers according to region or ethnicity, we conducted a linear regression analysis and explored age-by-sex interactions to determine the statistical significance of the nonparallel pattern in each cohort (see methods). Our analysis revealed that nonparallel patterns were more common in stomach cancer, melanoma, and thyroid cancer (Fig. 1B).

### Sex bias in tissue environments

A potential explanation for the prevalence of nonparallel patterns is that tissue environments in the stomach, skin, and thyroid exhibit greater sex differences compared with tissue environments in the other regions; these differences may influence carcinogenesis. To test this hypothesis, we used GTEx data to evaluate sex bias in tissue environments via sex-biased expressed pathway analysis. In our analysis, the level of expression in a pathway was represented by the mean level of expression for all genes in that pathway; we randomly selected 100 samples from each tissue to maintain consistent statistical power across tissues. We explored sex-biased expressed pathway using Limma (21); we repeated this process 1000 times and recorded the numbers of sex-biased expressed pathways in each tissue during each repetition (see methods). There were fewer than 100 kidney and bladder samples (Table S1) and no significant sex-biased expressed pathways were identified in those tissues; thus, their results were regarded as not available (NA). The results showed that the adrenal glands, stomach, thyroid, and skin contained the largest numbers of sex-biased expressed pathways (Fig. 2A).

**Fig. 2:**
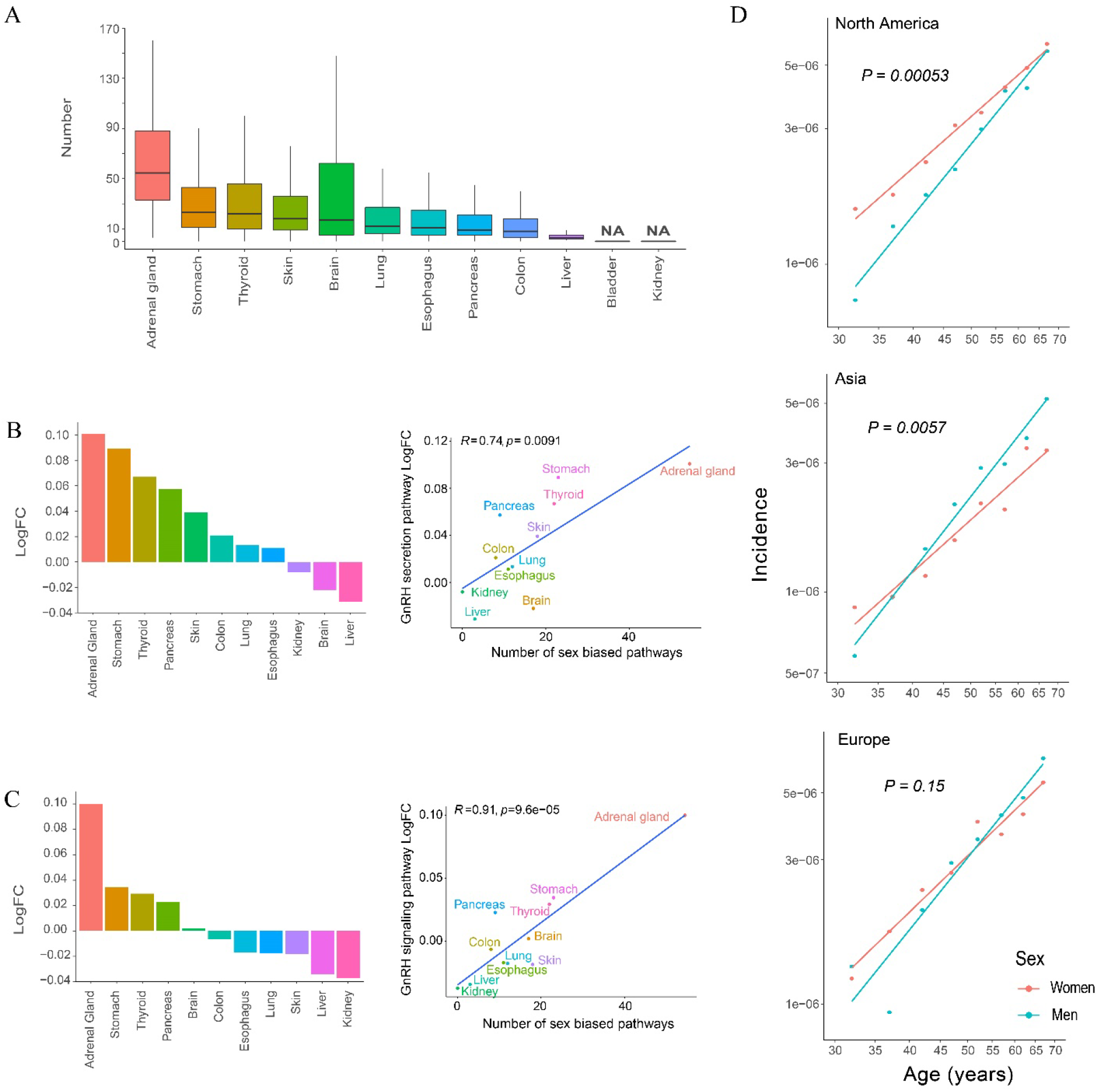
Sex bias in tissue environments and nonparallel patterns in adrenal gland cancer. (A) Number of sex-biased pathways expressed in each tissue. The results of bladder and kidney analyses were regarded as not available (NA) because of small sample size. (B and C) Female-to-male log fold changes of GnRH secretion and GnRH signaling pathways among tissues. The results of bladder analyses were excluded because of small sample size. (D) Nonparallel patterns in adrenal gland cancer. Regional cohorts within a single continent were merged to form a continental cohort.

To determine which pathways have greater involvement in tissue sex bias, we re-examined sex-biased expressed pathways using all samples in each type of tissue except the bladder, for which only nine samples were available. Among the pathways, only the gonadotropin-releasing hormone (GnRH) secretion pathway (hsa04929) exhibited significant sex bias (Limma P < 0.05) in the top three tissues with sex bias (adrenal glands, thyroid, and stomach), but not in other tissues (P for skin = 0.0683, Table S2). Additionally, the female-to-male log fold change (LogFC) of the GnRH secretion pathway were highest in the adrenal glands, stomach, and thyroid; the LogFC in skin was fifth highest (Fig. 2B). The LogFC of the GnRH secretion pathway was positively correlated (Pearson correlation coefficient *R* = 0.74) with the number of sex-biased expressed pathways in each tissue. The LogFC of another GnRH-related pathway, the GnRH signaling pathway (hsa04912), was also positively correlated (Pearson correlation coefficient *R* = 0.91) with the number of sex-biased expressed pathways in each tissue. The adrenal glands, stomach, and thyroid had the highest LogFCs, whereas the skin had a negative LogFC (Fig. 2C, Table S3).

Adrenal gland cancer is a rare type of cancer, and linear representation of its age-specific incidence is hindered by noise. Therefore, it was not included in the nonparallel analysis shown in Figure 1B. However, because adrenal gland tissue had the highest number of sex-biased expressed pathways and largest LogFC of the two GnRH-related pathways, we repeated the nonparallel pattern analysis with a focus on adrenal gland cancer. To reduce noise in the incidence of adrenal gland cancer, we merged the regional cohorts into continental cohorts (see methods). We found that adrenal cancer incidence also displayed a nonparallel pattern; the results were statistically significant in North America and Asia cohorts (Fig. 2D).

The most prevalent nonparallel patterns in stomach cancer, melanoma, thyroid cancer, and adrenal cancer were observed across regions or ethnicities (Figs. 1B and 2D). Consistent with these findings, adrenal glands, stomach, thyroid, and skin exhibited the highest numbers of sex-biased expressed pathways (Fig. 2A), which were associated with GnRH-related pathways. Thus, sex-biased tissue environments may influence the evolutionary dynamics of carcinogenesis and induce nonparallel patterns.

### Genes with sex-biased mutation frequency

Because carcinogenesis can be influenced by sex, there may exist genes that exhibit a greater likelihood of mutation in a single sex (hereafter, “sex-biased mutated genes”). Previous studies attempted to explore this hypothesis, but they had less statistical power because of reliance on incomplete The Cancer Genome Atlas (TCGA) data (16, 17) or whole genome sequencing data alone (18). Therefore, we used complete TCGA data to conduct a comparative analysis of gene mutation frequency between women and men among 6181 individuals across 24 non-reproductive cancers; we utilized both the likelihood ratio test (LRT) and logistic regression analysis to adjust for confounding factors (see methods). Our results revealed eight sex-biased mutated genes (Table 1): *DNAH7*, *LARP4B*, *PGM5*, and *PTEN* in stomach adenocarcinoma (STAD); *SELP* and *SPAG17* in skin cutaneous melanoma (SKCM); *RBM10* in lung adenocarcinoma (LUAD); and *CASP8* in head and neck squamous cell carcinoma (HNSC).

**Table 1.** Sex-biased mutated genes.

| Gene | Chromosome | Cancer | Female samples +/- | Male samples +/- | Relative risk (f/m) | LRT P value |
| --- | --- | --- | --- | --- | --- | --- |
| *CASP8* | 2 | HNSC | 23/108 | 19/332 | 3.24 | 3.85e-10 |
| *RBM10* | X | LUAD | 9/255 | 18/205 | 0.42 | 3.52e-4 |
| *SELP* | 1 | SKCM | 13/149 | 8/249 | 2.58 | 7.71e-3 |
| *SPAG17* | 1 |  | 17/145 | 13/244 | 2.07 | 7.71e-3 |
| *DNAH7* | 2 | STAD | 13/131 | 11/259 | 2.21 | 4.70e-2 |
| *LARP4B* | 10 |  | 13/131 | 8/262 | 3.05 | 4.27e-2 |
| *PGM5* | 9 |  | 13/131 | 8/262 | 3.05 | 4.27e-2 |
| *PTEN* | 10 |  | 14/130 | 12/258 | 2.19 | 4.70e-2 |

"Female samples +/-" and "Male samples +/-" refer to the numbers of samples from women or men that did or did not include a sex-biased gene mutation. "Relative risk (f/m)" represents the female-to-male relative risk of gene mutation frequency. "LRT P value" is the P value obtained from the likelihood ratio test, adjusted using the Benjamini-Hochberg method.

### Hotspot dominance in three sex-biased mutated genes in STAD

In STAD, three sex-biased mutated genes (*PTEN*, *LARP4B*, and *PGM5*) displayed hotspot mutation dominance, as shown in Figure 3 [depicted using ProteinPaint (22)]. These hotspots, Lys267ArgfsTer9 (c.800delA) in *PTEN*, Thr163HisfsTer47 (c.487delA) in *LARP4B*, and Ile98Val (c.292A>G) in *PGM5*, also showed a significant sex bias (P < 0.05, Fisher’s exact test, Fig. 3). TCGA data showed that these three hotspots were exclusively present in STAD, colon adenocarcinoma (COAD), and uterine corpus endometrial carcinoma (UCEC), with the exception of a *PGM5* hotpot in prostate adenocarcinoma (PRAD). The mutational spectra of other sex-biased mutated genes are shown in Figure S1.

**Fig. 3:**
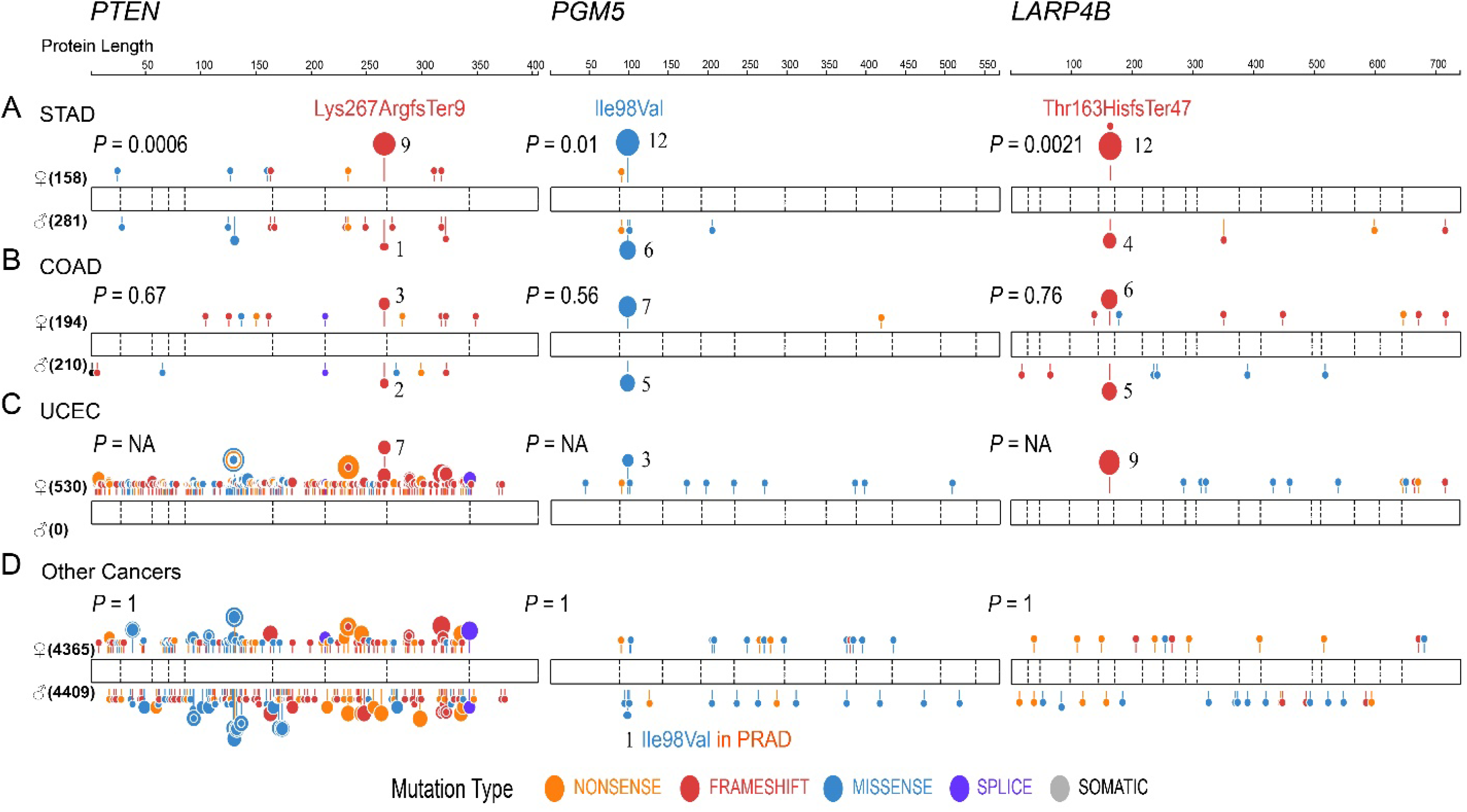
Mutational spectra of *PTEN*, *PGM5*, and *LARP4B*. P values above gene body on the left denote statistical significance of hotspot sex bias. Because UCEC is only present in women, the P value is regarded as NA. The numbers of female and male samples for each cancer are indicated on the left in each row. Mutation types are shown in different colors and are annotated at the bottom of the figure.

### Sex-biased hotspots and sex-biased proportions of cancers with microsatellite instability (MSI)

To determine the underlying cause of sex bias in the presence of the three hotspots, we conducted co-occurrence and mutual exclusivity analyses of the three hotspots in STAD, COAD, and UCEC; the results suggested that the three sex-biased hotspots tend to co-occur (Table S4). To gain further insight, we divided the samples into two groups (HOTSPOT+ and HOTSPOT-) according to the presence or absence of at least one of the three hotspots across STAD, COAD, and UCEC. Subsequently, we used Limma to conduct an analysis of differentially expressed genes (DEGs) between the two groups.

In all three cancers, four genes were differentially expressed between the two groups: *C7orf31*, *MLH1*, *EPM2AIP1*, and *LINC00667* (Fig. S2A, Data S4). *MLH1*, a well-known tumor suppressor gene, showed reduced expression in the HOTSPOT+ group across all three cancers (Fig. S2B). *MLH1* deficiency can induce MSI, which is commonly observed in colorectal, gastric, and endometrial cancer. The signatures of the three hotspots were consistent with previously reported mutational signatures of MSI cancer types: the *PTEN* and *LARP4B* hotspots are typical of the MSI mutational signature ID1, whereas the *PGM5* hotspot is typical of the MSI mutational signatures SBS21 and SBS26 (23). Furthermore, all HOTSPOT+ samples were labeled as the MSI subtype in TCGA data (see methods). This evidence suggests that the three hotspots result from hypermutation across MSI subtypes.

We speculated that hotspot-specific sex bias is caused by sex bias in MSI subtype. Thus, we examined MSI subtype proportions in STAD and COAD; in both cancers, we found that the MSI subtype proportion was higher in women than in men. The difference was statistically significant in STAD (Table S5). A previous study (24) revealed that, in both gastric and colorectal cancers, the MSI subtype proportion was significantly higher in women than in men (Table S6).

### Estrogen receptor alpha (*ESR1*), thyroid hormone receptors (THRs), and hypoxia-inducible factors (HIFs) in cancers with MSI

The above findings led us to investigate the underlying cause of sex bias in MSI subtype. For this purpose, we divided STAD, COAD, and UCEC samples into two groups: MSI-positive (MSI+) and MSI-negative (MSI-). We then analyzed DEGs in each cancer type (see methods).

Hypoxia is known to suppress mismatch repair through mechanisms such as the reduction of *MLH1* expression (25–28). In STAD, COAD, and UCEC, we observed that MSI+ samples consistently had a significantly elevated level of *HIF1A* (a key activator of hypoxia signaling) or a reduced level of *HIF3A* [a repressor of hypoxia signaling (29)], compared with MSI-samples. These findings indicated that MSI+ samples had a higher level of hypoxia. Subsequently, we used the ratio of *HIF1A* to *HIF3A* (i.e., HIF level), to represent the level of hypoxia in each sample. We found that the HIF level was significantly higher in MSI+ samples than in MSI-samples; it was positively correlated with the *MLH1* methylation level in all three cancers (Fig. 4A).

**Fig. 4:**
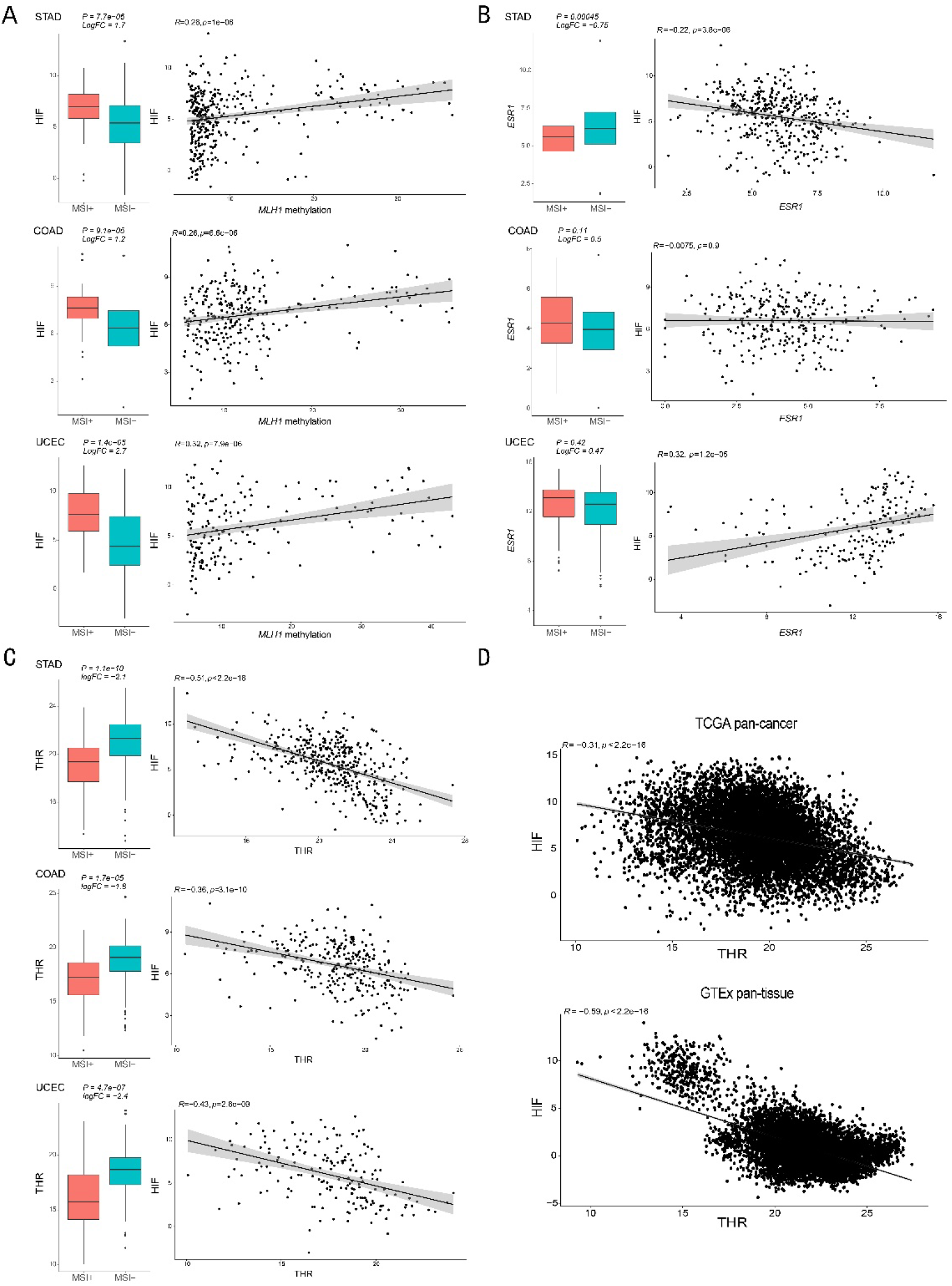
Correlations among HIF, *ESR1*, and THR. (A) Comparison of HIF levels between MSI+ and MSI-samples, and correlations between HIF level and *MLH1* methylation level across STAD, COAD, and UCEC. (B) Comparison of *ESR1* levels between MSI+ and MSI-samples, and correlations between HIF level and *ESR1* level across STAD, COAD, and UCEC. (C) Comparison of THR levels between MSI+ and MSI-samples, and correlations between HIF level and THR level across STAD, COAD, and UCEC. (D) Correlations between HIF level and THR level in pan-cancer and pan-tissue analyses. In all dot plots, *R* represents the Pearson correlation coefficient; P values were obtained using Student’s t-test.

*ESR1* has been reported that it can directly regulates *HIF1A*, indicating crosstalk between the hypoxia and estrogen signaling pathways (30). However, in STAD, COAD, and UCEC, we found that the level of *ESR1* did not significantly differ between MSI+ and MSI-samples. Furthermore, the expression of *ESR1* was positively correlated with the HIF level in UCEC, but not in STAD or COAD (Figs. 4B and 5). Finally, pan-cancer analysis of TCGA data showed that *ESR1* only had a slight positive correlation with the HIF level; conversely, pan-tissue analysis of GTEx data revealed a negative correlation with the HIF level (Fig. S3).

**Fig. 5:**
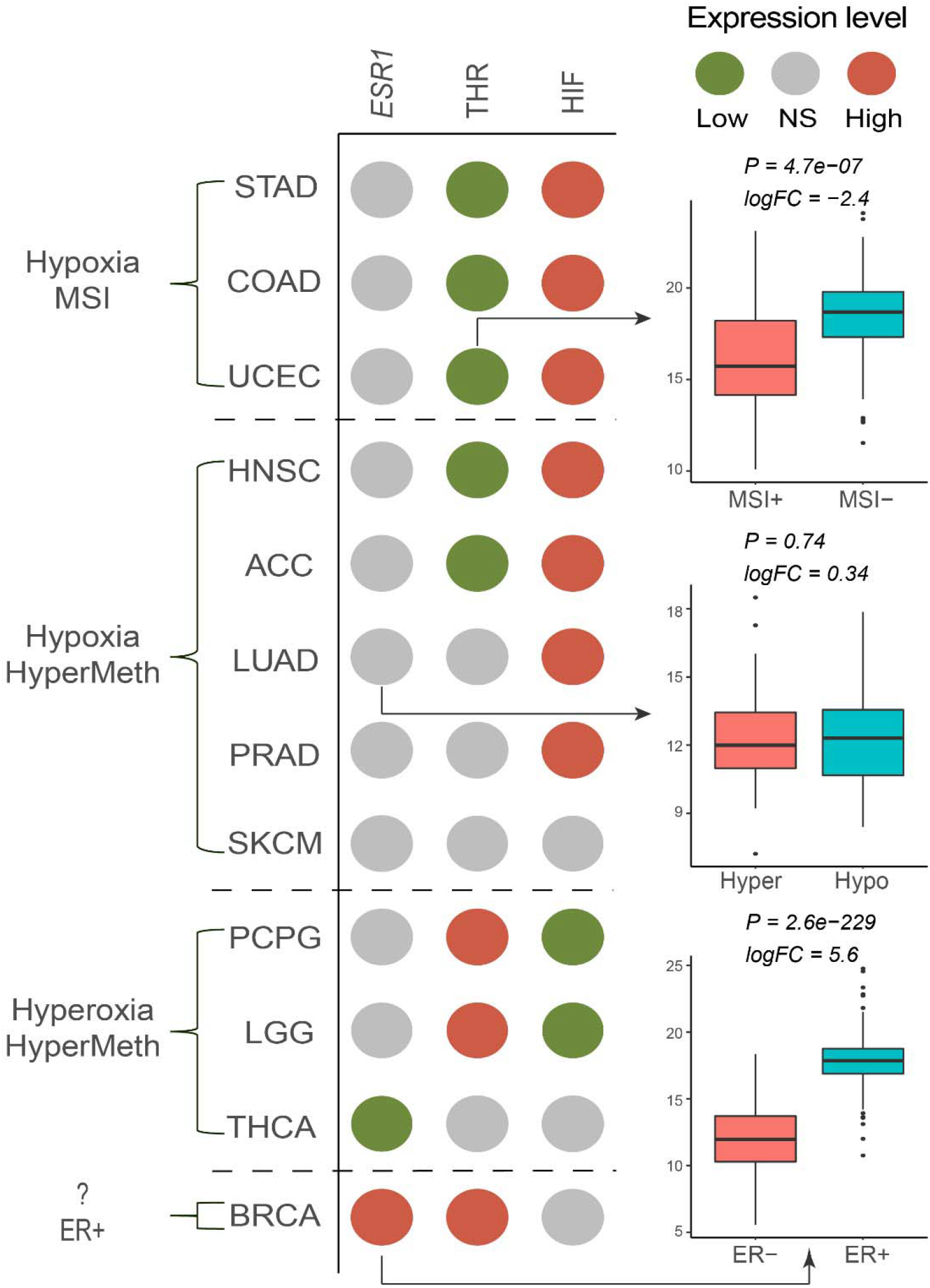
HIF, *ESR1*, and THR expression signatures in TCGA cancers. Comparative expression signatures of *ESR1*, THR, and HIF are represented by colored dots. Red dots indicate significantly elevated expression, green dots indicate significantly reduced expression, and gray dots indicate no significant difference in expression when MSI+, HyperMeth, and ER+ cancers are compared with control cancers. Boxplots on the right are representative examples to illustrate the meaning of the dots. LGG: low-grade glioma, THCA: thyroid carcinoma.

Our above results indicated that GnRH was involved in the establishment of a sex-biased tissue environment. GnRH is a key regulator of the hypothalamic-pituitary-gonadal (HPG) axis, which is upstream of estrogen signaling and has substantial crosstalk with the hypothalamic-pituitary-adrenal (HPA) and hypothalamic-pituitary-thyroid (HPT) axes (31). Considering these connections, besides *ESR1*, we suspected that other hormones or hormone receptors were involved in the sex bias in MSI subtype. Therefore, we investigated the expression patterns of a set of genes involved in the HPG, HPA, and HPT axes (see methods).

The results showed that only one gene, thyroid hormone receptor beta (*THRB*), was significantly differentially expressed between MSI+ and MSI-samples in all three cancer types. Moreover, thyroid hormone receptor alpha (*THRA*) was significantly differentially expressed in STAD and COAD, but not in UCEC (Fig. S4, Data S5). Because *THRA* and *THRB* are both important receptors for thyroid hormone (TH), we used the summed logarithmic levels of *THRA* and *THRB* to represent the THR level in each sample. We found that the THR level was significantly lower in MSI+ samples than in MSI-samples; it was negatively correlated with the HIF level in all three cancers (Fig. 4C). The negative correlation between HIF and THR was also confirmed by pan-cancer analysis of TCGA data and pan-tissue analysis of GTEx data (Fig. 4D).

### *ESR1*, THR, and HIF in cancers with hypermethylation or in estrogen receptor-positive (ER+) breast cancers (BRCAs)

Epigenetic silencing of the *MLH1* gene is a major cause of MSI; it can occur in cancers with the CpG island methylator phenotype (CIMP) or hypermethylation. There is a significant overlap between colorectal cancers with MSI and colorectal cancers with CIMP (32). Accordingly, to explore whether cancers with CIMP or hypermethylation exhibit expression signatures similar to MSI+ cancers (e.g., *ESR1*, THR, and HIF), we conducted a similar analysis of cancers with sufficient CIMP or hypermethylation labels in the TCGA dataset. These cancers were designated “HyperMeth cancers,” whereas other cancers were designated “HypoMeth cancers” (see methods).

Based on their expression signatures, HyperMeth cancers were divided into two subgroups: hypoxia and hyperoxia. In the hypoxia subgroup, similar to MSI+ cancers, all HyperMeth cancers demonstrated a significantly elevated HIF level (except HyperMeth SKCM, in which the increase was not statistically significant); they typically had a reduced THR level or an elevated *ESR1* level (Figs. 5 and S6). In the hyperoxia subgroup, all HyperMeth cancers demonstrated a significantly reduced HIF level (except HyperMeth THCA, in which the decrease was not statistically significant); they also had a significantly elevated THR level or a significantly reduced *ESR1* level (Fig. 5).

Adrenocortical carcinoma (ACC) and pheochromocytoma and paraganglioma (PCPG) both originate from adrenal glands. However, HyperMeth ACC and HyperMeth PCPG displayed opposing THR and HIF expression signatures. Specifically, HyperMeth ACC exhibited hypoxia, whereas HyperMeth PCPG exhibited hyperoxia. This difference suggests that the two cancers originated under divergent selective pressures or evolved in opposite directions.

BRCA is classified into four well-characterized molecular subtypes: luminal A, luminal B, HER2-positive, and triple-negative. Estrogen plays a crucial role in the development of ER+ BRCAs (i.e., luminal A and luminal B). To investigate whether ER+ BRCAs exhibit hypoxia and have expression signatures similar to MSI+ or hypoxic HyperMeth cancers, we compared expression signatures between ER+ BRCAs and ER-(i.e., HER2-positive and triple-negative) BRCAs. In contrast to the findings in MSI+ and HyperMeth cancers, we observed an antagonistic expression pattern in ER+ BRCAs: *ESR1* and THR levels were both significantly elevated, whereas the HIF level did not significantly differ (Fig. 5). However, the *HIF1A* level tended to be significantly reduced, and the *HIF3A* level was significantly reduced (Fig. S5).

Estrogen receptor beta (*ESR2*) is also involved in hypoxia signaling. However, its role is not well established, and it exhibits opposing roles in some cancers (33). Our results showed that *ESR2* was significantly elevated in MSI+ COAD and significantly reduced in HyperMeth SKCM, indicating that *ESR2* may play a role in the initiation of these cancers.

Additionally, similar to the three hotspot mutations exclusively found in MSI+ cancers, some other sex-biased mutated genes tend to be present in certain cancer subtypes. Specifically, *CASP8* was typically mutated in HyperMeth HNSC, *SELP* was typically mutated in HyperMeth SKCM, and *DNAH7* was typically mutated in MSI+ STAD. These results were statistically significant (Table S7), suggesting a link between these sex-biased mutated genes and the corresponding cancer subtypes.

## Discussion

The above results imply that a sex-biased tissue environment can influence carcinogenesis. *ESR1* and THR are associated with the HIF level in cancers, as well as specific cancer subtypes, suggesting that they also play a role in carcinogenesis. *ESR1* can directly regulate *HIF1A* and induce the hypoxia signaling pathway (30). TH has a more complex role in hypoxia. TH can induce *HIF1A* activity, which then reduces TH signaling; however, experimental and clinical studies have revealed that TH plays a critical role in protecting tissues from hypoxic injury (34). We also observed this complex phenomenon: analysis of GTEx data revealed that the THR level was negatively correlated with the *HIF1A* level but positively correlated with the *HIF3A* level (Fig. S7). However, analysis of TCGA data demonstrated that the negative correlation between THR and *HIF1A* disappeared, whereas the positive correlation between THR and *HIF3A* persisted (Fig. S8). These results indicated that THR generally functions as a hypoxia repressor in cancers, rather than a hypoxia activator, similar to its protection against hypoxic injury. ERs and THRs are both nuclear receptors with similar DNA-binding sequences. They can competitively bind to each other’s DNA-binding sites and inhibit subsequent transcription (35). The mutual antagonism of ERs and THRs is consistent with the observations in Figure 5: an increase in *ESR1* or a decrease in THR is accompanied by hypoxia, whereas a decrease in *ESR1* or an increase in THR is accompanied by hyperoxia. Based on the present findings, we propose a model of carcinogenesis that involves imbalances in estrogen and TH (Fig. 6). As noted above, TH and estrogen play key roles in many aspects of biological processes and maintain homeostasis under healthy conditions. However, TH or estrogen dominance can induce changes in the tissue environment, such as hypoxia or hyperoxia; it can also alter the epigenome and transcriptome in tissue cells. In ER+ BRCAs, both TH and estrogen can become simultaneously elevated, resulting in a state of antagonism or “chaotic-oxia.” In terms of selective pressure for cancer evolution, all of these altered tissue environments can induce specific cancer subtypes, as shown in Figure 5 and 6.

**Fig. 6:**
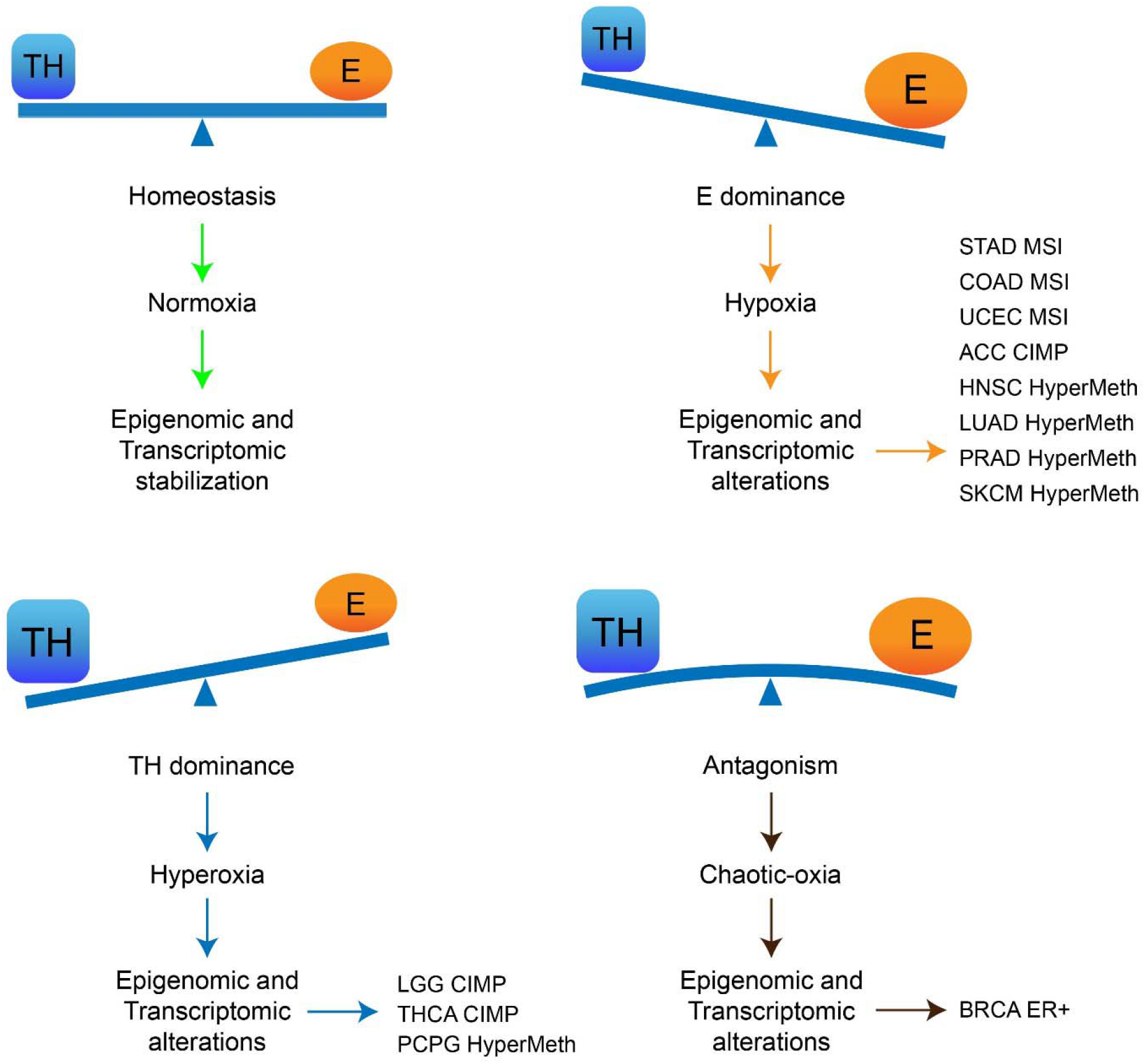
The potential contributions of imbalances in estrogen and TH to carcinogenesis. The relationship between estrogen (E) and TH is illustrated by a seesaw diagram. In a state of homeostasis, the levels of E and TH are in balance. When this balance is disrupted, epigenomic and transcriptomic alterations can occur in affected tissues. These alterations can lead to changes in the tissue environment and function as a selective pressure, thereby inducing specific cancer subtypes.

The present study showed that carcinogenesis may be influenced by sex bias in two ways: sex-biased tissue environments and hormonal imbalances. However, these influences are not mutually exclusive and may interact with each other. Here, we found that GnRH was associated with a sex-biased tissue environment; GnRH is a key regulator of the HPG axis and receives negative feedback from gonadal steroids, such as estrogen (36). The HPG axis has extensive crosstalk with the HPT axis, such as crosstalk between the estrogen and TH signaling pathways (31, 35). Together, the HPG axis and HPT axis, including estrogen and thyroid hormones, regulate tissue environment homeostasis. Generally, the level of estrogen is higher in women than in men (37); this increases the likelihood of a hypoxic tissue environment in women and the potential for cancers with MSI. As expected, we found that women had a higher prevalence of MSI in STAD and COAD; we also found that women had a high prevalence of MSI in UCEC, where estrogen is suspected to promote cancer progression (Table S5 and S6). In contrast, the lower level of estrogen in men increases the likelihood of a hyperoxic tissue environment in men and the potential for cancers such as the hyperoxic CIMP subtype of thyroid carcinoma. Although the difference was not statistically significant (possibly because of the small sample size), we found that men had a higher prevalence of the hyperoxic CIMP subtype of thyroid carcinoma (Table S5). However, we found that the hypoxic HyperMeth subtype of HNSC was more prevalent in men, indicating that it may exhibit tissue-specific regulation. These results suggest that in the context of sex-biased tissue environments and hormonal imbalances, cancer precursors have a preferred evolutionary trajectory. Consistent with this hypothesis, we observed that six of the eight sex-biased mutated genes were significantly enriched in specific cancer subtypes (Table S7). However, in cancers such as PCPG and SKCM, the proportions of subtypes were similar in women and men.

In addition to influencing the direction of cancer evolution, sex-biased tissue environments and hormonal imbalances may also affect the evolutional pace of carcinogenesis. GnRH secretion exhibits pulsatile behavior in both men and women, but the frequency and amplitude of pulses in women are dependent on the phase of the menstrual cycle which do not exist in men (36). The sex-biased dynamics of GnRH secretion may lead to sex-biased dynamics of cancer evolution. Hypermutation or hypermethylation, which occur in cancers with hormonal imbalances, could lead to more rapid cancer evolution and altered dynamics of carcinogenesis, as described in a previous study (6). All of these factors may contribute to the nonparallel pattern of cancer incidence observed in this study. However, greater understanding of sex-biased biological mechanisms is needed to develop a detailed mathematical model to explain the sex bias underlying the two patterns of age-specific cancer incidence.

TH is suspected to play a role in carcinogenesis. For example, compared with men, women are 10-fold more likely to develop hyperthyroidism and hypothyroidism (38); these disorders have been linked to changes in the incidence and prognosis of many cancers (39). The present findings support the increasing importance of TH in cancer research. Although our study focused on the balance between TH and estrogen, we acknowledge that other hormones may also contribute to carcinogenesis. For example, progestogen is reportedly involved in luminal A BRCA, and androgens have been linked to prostate cancer.

In this study, samples were grouped by molecular clustering labels, which may not fully capture the status of the balance between estrogen and TH. The identification of molecular patterns associated with imbalances in estrogen and TH could enhance our findings and reveal additional mechanisms involving sex bias. Furthermore, the levels of ERs and THRs may not directly reflect the levels of estrogen and TH. Nevertheless, this study showed that three phases (hypoxia, hyperoxia, and chaotic-oxia) are present in most cancers and are linked to hormonal imbalances.

## Materials and methods

### Epidemiology data processing

To ensure the reliability of our analysis, we selected regions with the top 10 GDP in 2005, corresponding to the CI5X database, which records incidence from 2003 to 2007. The GDP information was obtained from the World Bank database (https://databank.worldbank.org/). In the data of USA, we selected two cohorts – white and black – to cover the black race, resulting in a total of 11 cohorts of regions or races: USA black, USA white, Japan, Germany, England, China, France, Italy, Canada, Spain, and Korea. We then chose major non-reproductive cancers according to Cancer Statistics US (40): lung (LUSC and LUAD), melanoma, colon, bladder, kidney, pancreas, thyroid, rectum, liver, stomach, brain, esophagus, oral cavity and pharynx, gallbladder, and larynx. This resulted in a total of 11 by 16 cohorts included in our analysis.

To reduce noise in the incidence of adrenal gland cancer, we merged the regional cohorts into continent cohorts. Specifically, we merged the USA black, USA white, and Canada cohorts into a North America cohort, merged Japan, China, and Korea cohorts into an Asia cohort, and merged Germany, England, France, Italy, and Spain cohorts into a Europe cohort.

We cut the age-specific incidence data from 30 to 70 years of age. Because previous studies have shown that incidence rates before the age of 30 are higher than expected due to factors such as family inheritance, exposure to carcinogens, or explosive cell division during development (1, 41), and incidence rates after the age of 70 are lower than expected due to the selection of relatively unsusceptible individuals (42). We then used the least squares method to fit straight lines separately for women and men at a log-log scale.

### Nonparallel pattern test

We consider a linear equation (equation 1) as below:

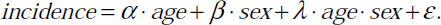

Denoting 0 as female and 1 as male, so the linear equation for male is:

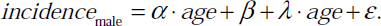

The linear equation for female is:

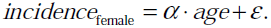

When λ is 0, two lines of sexes are parallel, when λ is not 0, two lines of sexes are nonparallel. So, we perform a linear regression as equation 1, and test if λ is significant not equal to 0 (significant nonparallel, Student’s t-test). Then P values are adjusted by Bonferroni methods, adjusted P values which lower that 0.05 are reported as significant.

### Sex-biased expressed pathways analysis

Carcinomas typically originate from solid tissues. Therefore, in the GTEX data, we selected tissues from which major solid cancers originate and excluded blood and solid tissues that are less likely to develop cancer, such as heart and fat. After this selection process, we chose 12 tissues: bladder, kidney, liver, skin, colon, stomach, pancreas, lung, thyroid, brain, esophagus, and adrenal gland. The number of samples for each tissue is shown in Table S1.

We obtained human pathway information using the R package KEGGREST (43, 44), resulting in a total of 352 pathways included in our analysis.

To ensure that each tissue had equal testing power for comparison between tissues, we randomly selected 100 samples from each tissue and conducted sex-biased expressed pathways analysis. For each pathway, we used the mean logarithmic expression [log (counts+1)] of all genes in the pathway as the expression strength of the pathway. We divided the samples in each tissue into two groups based on sex and compared pathway expression strength between groups using Limma, while correcting for the factor of the position of the tissue (body site detail). As shown in Table S1, there was a large difference in sample size between tissues, and our approach allowed us to account for these differences and ensure that our analysis was statistically robust.

Our analysis was affected by poor testing power due to both the expression strength of the pathway, the mean expression of all genes, can weaken the real signal of a set of specific genes in that pathway, and the small sample size resulting from our sampling approach. However, as we only made qualitative comparisons, we relaxed the significance standard and considered unadjusted P values < 0.05 to be significant. We then recorded the number of significantly different pathways in each tissue. This analysis process was repeated 1000 times, and the numbers of sex-biased expressed pathways in each tissue were displayed in Figure 2.1C. As the samples from bladder and kidney were less than 100, and they had no significant sex-biased expressed pathways, their final statistical results were marked as NA.

### Mutation data processing for sex-biased mutated gene test

We used MC3 profile of TCGA mutation data. We excluded pediatric and blood cancers, as well as sex-specific cancers, leaving only non-reproductive adult solid tumors.

To filter out passenger mutations and reduce data noise, we performed filtering on the mutation data. Firstly, we retained mutations of the following types: nonsense, insertion, deletion, translation start site, and non-stop mutations, as these mutations would theoretically change the protein sequence. We also retained mutations predicted as “deleterious” by SIFT (45) and as “probably damaging” by PolyPhen (46). In addition, we retained mutations at a single site where mutations occurred five or more times in the entire MC3 dataset.

To reduce potential statistical bias from extremely high mutation samples, we removed the top 5% of samples with the highest tumor mutation burden (TMB) for each cancer type. Additionally, we removed genes with a sample mutation rate below 5%, as these genes have limited biological significance and their removal can improve the power of a statistical test. For the analysis of melanoma, which is only susceptible in white populations and typically only has nonparallel patterns in white populations, we only considered the white population.

### Likelihood ratio test

Based on our preliminary analysis of sex-biased mutated genes using the Fisher exact test and the conclusions from previous work (47), we identified tumor mutation burden (TMB) as an important confounding factor for testing sex-biased mutated genes, as there are sex-biased TMB in many cancers. Failure to exclude this confounding factor can result in many false positives, particularly for super long genes. To address this issue, we used a likelihood ratio test (LRT) similar to previous work (17) to eliminate the confounding factor of TMB. The LRT also corrected for the difference in copy numbers of the X chromosome between men and women.

The background mutation rate varies between samples. Firstly, we assumed that the mutation rate of a gene between male and female samples is unbiased. Therefore, the mutation rate ratio of a gene between male and female samples should be approximately equal to the background mutation rate ratio between male and female samples. We formulated the null hypothesis that the mutation rate of a gene is not biased between male and female samples. In other words, if the null hypothesis is true, for each gene:

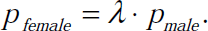

Where *P_female_*is the mutation rate of a gene in female samples, *P_male_* is the mutation rate of a gene in male samples, and λ is the background mutation rate ratio between male and female samples. We defined the mutation rate of a gene as the ratio of samples with mutations in that gene divided by the total number of female or male samples in a given cancer type. We denoted the mutation rate of a gene in female samples as *P_female_* and the mutation rate of the same gene in male samples as *P_male_*. The background mutation rate ratio for a gene was defined as the ratio of the total number of mutations on the chromosome where the gene is located in female samples to the total number of mutations on the same chromosome in male samples.

Under the assumption that the null hypothesis is true, the likelihood function is:

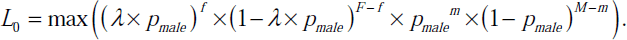

If the null hypothesis is not true, it is assumed that P_male_ and P_female_ are not proportional to each other, and the likelihood function is:

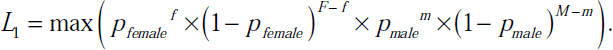

The log-likelihood ratio (LLR) is then calculated as:

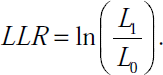

According to Wilks’ theorem, 2×LLR follows a chi-square distribution with one degree of freedom. Thus, we can calculate the P value, and the P value for each gene is adjusted using the Benjamini-Hochberg method within each cancer types.

### Logistic regression analysis

To account for potential confounding factors such as age, cancer grade, smoking history, and alcohol history, we conducted logistic regression analysis with these covariates to investigate the significance of sex in gene mutations.

We used the likelihood ratio test to identify genes with adjusted P values below 0.05, which helped eliminate the confounding factor of tumor mutation burden (TMB) and chromosome copy number. We also used logistic regression analysis with a significance threshold of P < 0.05 to identify genes that were significant after accounting for confounding factors. Genes that were identified as significant by both the likelihood ratio test and logistic regression analysis were reported as significant genes.

### Test for sex biased mutated genes on Y chromosome

As women do not possess the Y chromosome, the LRT method cannot be employed for genes located on the Y chromosome. Therefore, we utilized the Fisher exact test to evaluate the significance of mutations in genes located on the Y chromosome. We adjusted the resulting P values using the Benjamini-Hochberg method to account for multiple testing. Genes with an adjusted P value below 0.05 were considered significant.

### Subtype labeling in TCGA data and sample grouping

We obtained sample subtypes in TCGA from the molecular subtype profiles available on UCSC Xena (48). We used the following sample groupings for our analysis.

For STAD and COAD, we grouped samples labeled “HM-indel” or “HM-SNV” in the MSI+ group, while samples labeled “CIN”, “GS”, or “EBV” were grouped in the MSI-group. These sample groupings match well with the samples labeled “msi-h” in previous study (49). For UCEC, we grouped samples labeled “UCEC.POLE” or “UCEC.MSI” in the MSI+ group, while samples labeled “UCEC.CN_LOW” or “UCEC.CN_HIGH” were grouped in the MSI-group.

For LGG and ACC, samples labeled “CIMP-high” were grouped in the HyperMeth group, while samples labeled “CIMP-low” were grouped in the HypoMeth group. For THCA, samples labeled “CpG island methylated” were grouped in the HyperMeth group, while samples labeled “follicular” were grouped in the HypoMeth group. For SKCM and HNSC, samples labeled “CpG island-methylated” or “hyper-methylated” were grouped in the HyperMeth group, while samples labeled “normal-like” or “hypo-methylated” were grouped in the HypoMeth group. For PCPG, samples labeled “hyper-methylated” were grouped in the HyperMeth group, while samples labeled “low-methylated” were grouped in the HypoMeth group. For LUAD, samples labeled “high” were grouped in the HyperMeth group, while samples labeled “low” were grouped in the HypoMeth group. For PRAD, samples labeled “1” were grouped in the HyperMeth group, while samples labeled “4” were grouped in the HypoMeth group.

For BRCA, samples labeled “LumA” or “LumB” were grouped in the ER+ group, while samples labeled “Her2” or “Basal” were grouped in the ER-group.

### Differentially expressed genes (DEGs) analysis

We conducted DEGs analyses using Limma between two groups in each cancer type with enough labeled samples (more than 50). Specifically, we compared HOTSPOT+ versus HOTSPOT-in STAD, COAD, and UCEC; MSI+ versus MSI-in STAD, COAD, and UCEC; HyperMeth versus HypoMeth in HNSC, LUAD, PRAD, ACC, PCPG, LGG, THCA, and SKCM; and ER+ versus ER-in BRCA. We corrected for covariates such as sex, age, tumor stage, race, alcohol, and smoking history during the DEG analyses.

We created two pseudogenes, THR and HIF, to represent thyroid hormone receptor level and hypoxia level, respectively, in each sample. Pseudogene THR was created by using the summed logarithmic counts of *THRA* and *THRB*, while pseudogene HIF was created by using the logarithmic counts of *HIF1A* minus the logarithmic counts of *HIF3A*. Both THR and HIF were analyzed together with all the other genes during the DEG analyses.

We reported the adjusted P value and log fold change (logFC) provided by Limma for each gene. Specifically, genes with a Limma adjusted P value lower than 0.05 and an absolute value of logFC higher than 1 were considered significant.

### Sets of genes focused in expression signature

Basing on the role of GnRH on tissue sex bias and the crosstalk between HPG, HPA and HPT axes and relationship between *ESR1* and hypoxia, we focused on the following sets of genes: GnRH (*GNRH1*) and its receptors (*GNRHR*, *GNRHR2*), glycoprotein hormones alpha polypeptide (*CGA*), luteinizing hormone (*LHB*) and its receptors (*LHCGR*), follicle stimulating hormone (*FSHB*) and its receptor (*FSHR*), thyrotropin releasing hormone (*TRH*) and its receptor (*TRHR*), thyroid stimulating hormone (*TSHA*, *TSHB*) and its receptor (*TSHR*), corticotrophin-releasing hormone (*CRH*) and its receptor (*CRHR1* and *CRHR2*), adrenocorticotropic hormone (*POMC*) and its receptor (*MC2R*), glucocorticoid receptor (*NR3C1*), thyroxine-binding globulin (*SERPINA7*), transthyretin (*TTR*), estrogen receptors (*ESR1*, *ESR2*), thyroid hormone receptors (*THRA*, *THRB*), testis receptor (*NR2C1*, *NR2C2*), androgen receptor (*AR*), progesterone receptor (*PGR*), hypoxia-inducible factors (*HIF1A*, *EPAS1*, *HIF3A*, *EGLN1*, *EGLN2*, *EGLN3*, *ARNT*).

## Declarations

### Ethics approval and consent to participate

Research was carried out on data obtained from public data. No ethical approval was necessary as part of accessing these data.

### Consent for publication

Not applicable for this work.

### Data availability

The results published here are based on data generated by the TCGA Research Network (https://www.cancer.gov/tcga), the Genotype-Tissue Expression (GTEx) Project (https://www.gtexportal.org), UCSC Xena (https://xenabrowser.net/datapages), and the IARC database (http://ci5.iarc.fr/CI5-X/Pages/download.aspx). The analysis code is archived on GitHub (https://github.com/WhitePredator/sex-bias-in-cancer-initiation).

### Author contributions

F.Z. performed data processing, data analysis, interpreted the data, and prepared the manuscript.

M.D. acquired funding, supervised the study, and reviewed the manuscript.

## Acknowledgements

The authors thank Minping Qian, Hong Wu, Zexian Zeng, Ruibin Xi, Donglai Wang, Ge Gao, Qing Li, Chao Tang, Jinzhi Lei, Yihan Lin, and Zemin Zhang for insightful discussions.

## Supplementary Information

**Fig. S1.**
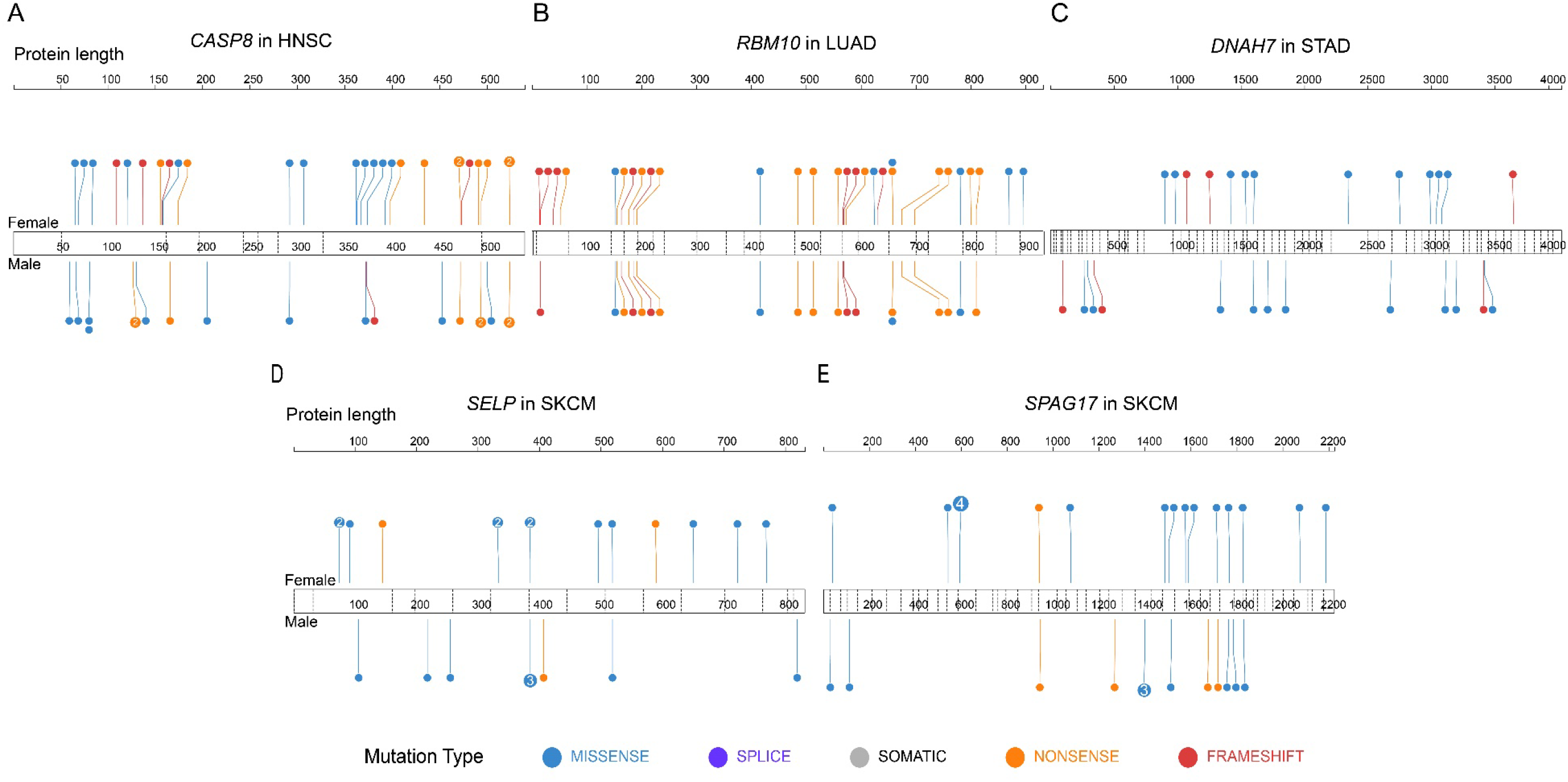
Mutational spectra of other sex-biased mutated genes in specific cancers.

**Fig. S2.**
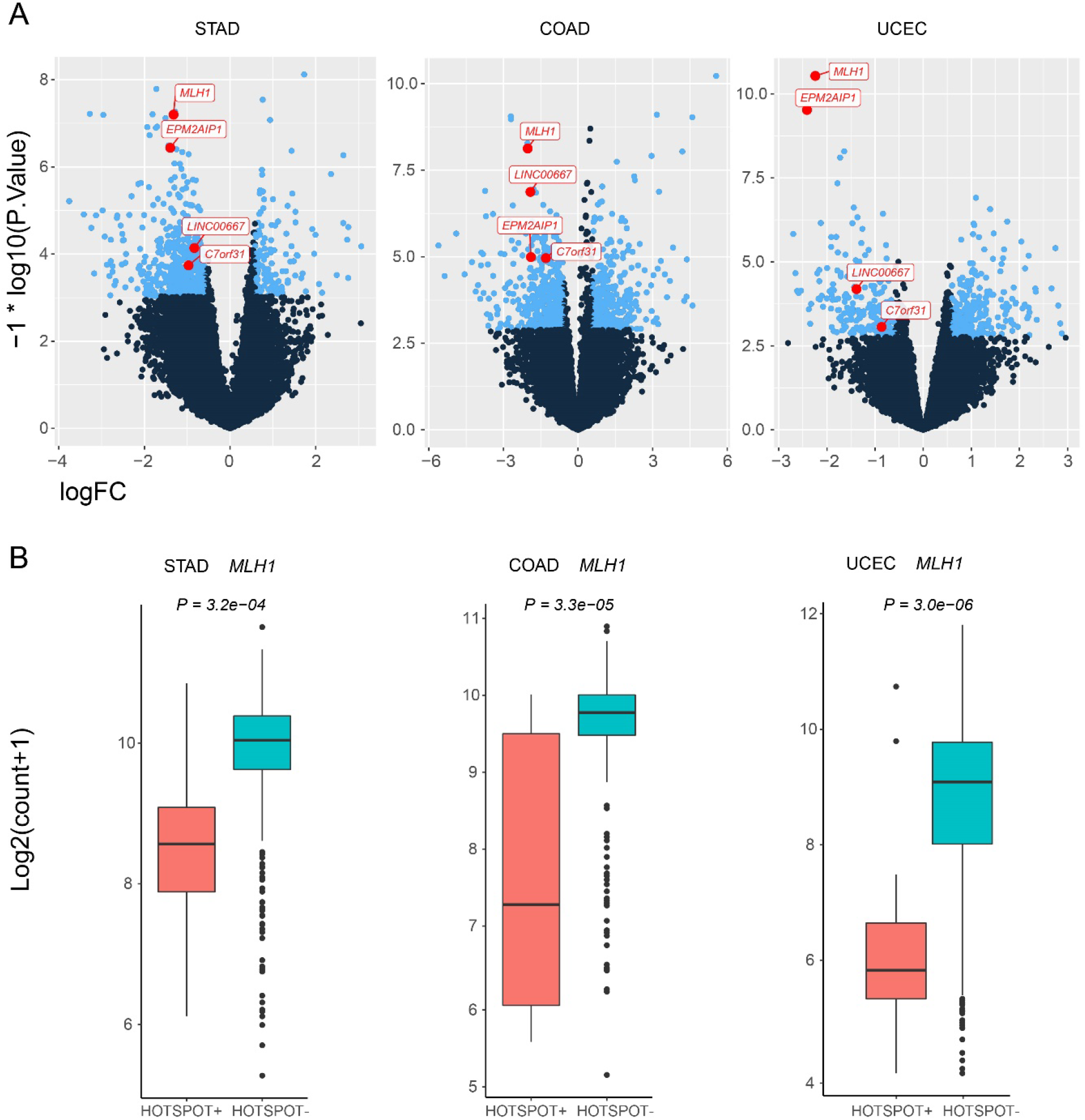
*MLH1* expression in HOTSPOT+ and HOTSPOT-groups. Analyses of DEGs were conducted between HOTSPOT+ and HOTSPOT-groups in STAD, COAD, and UCEC using Limma. The volcano plots are shown in figure A, and four significantly genes are denoted in red. *MLH1* expression is shown in the figure B. P values shown are adjusted P values obtained using Limma.

**Fig. S3.**
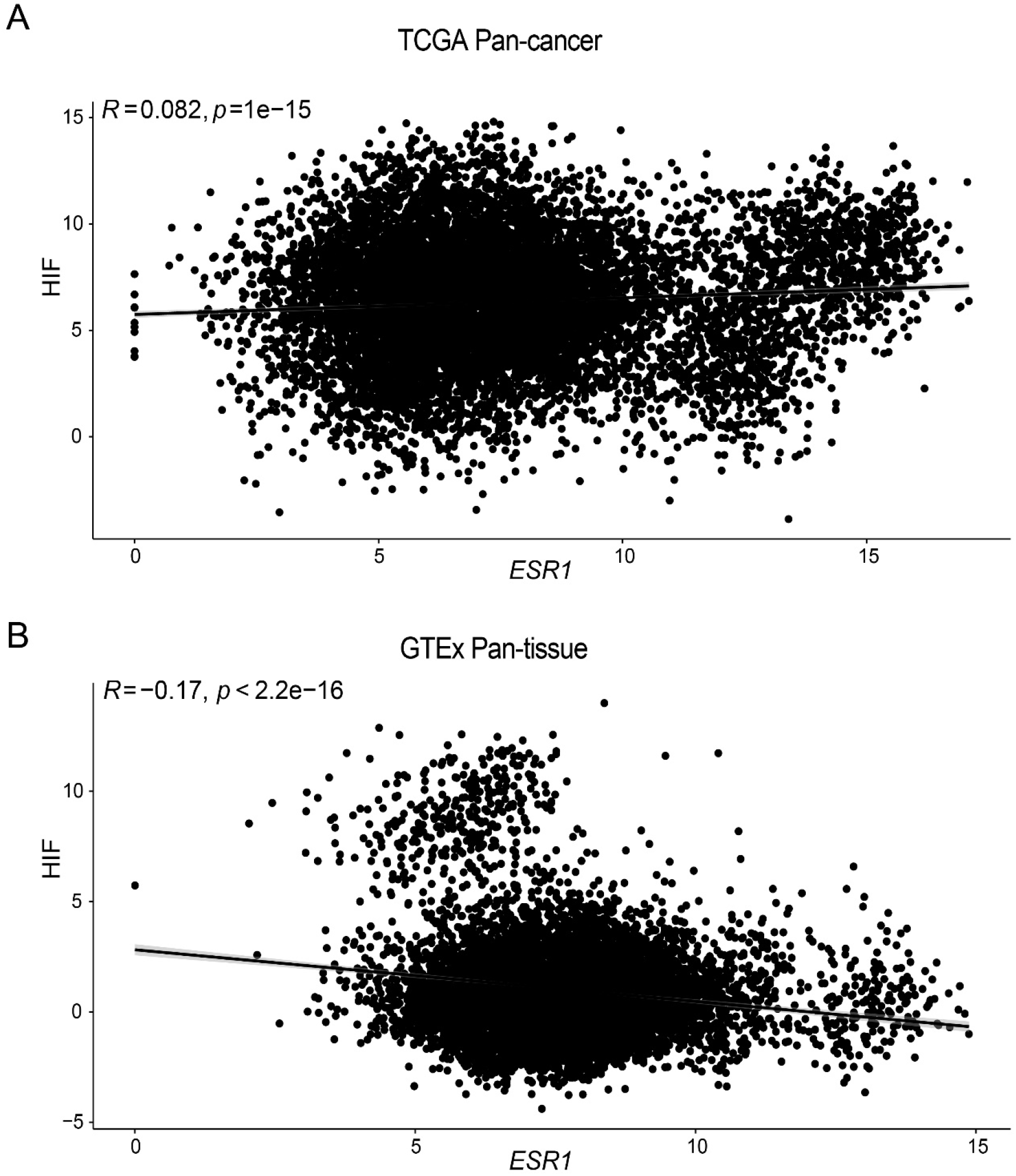
Correlations between HIF and *ESR1*. Correlations between HIF level and *ESR1* level in pan-cancer analysis of TCGA data and in pan-tissue analysis of GTEx data. *R* represents the Pearson correlation coefficient; P values were obtained using Student’s t-test.

**Fig. S4.**
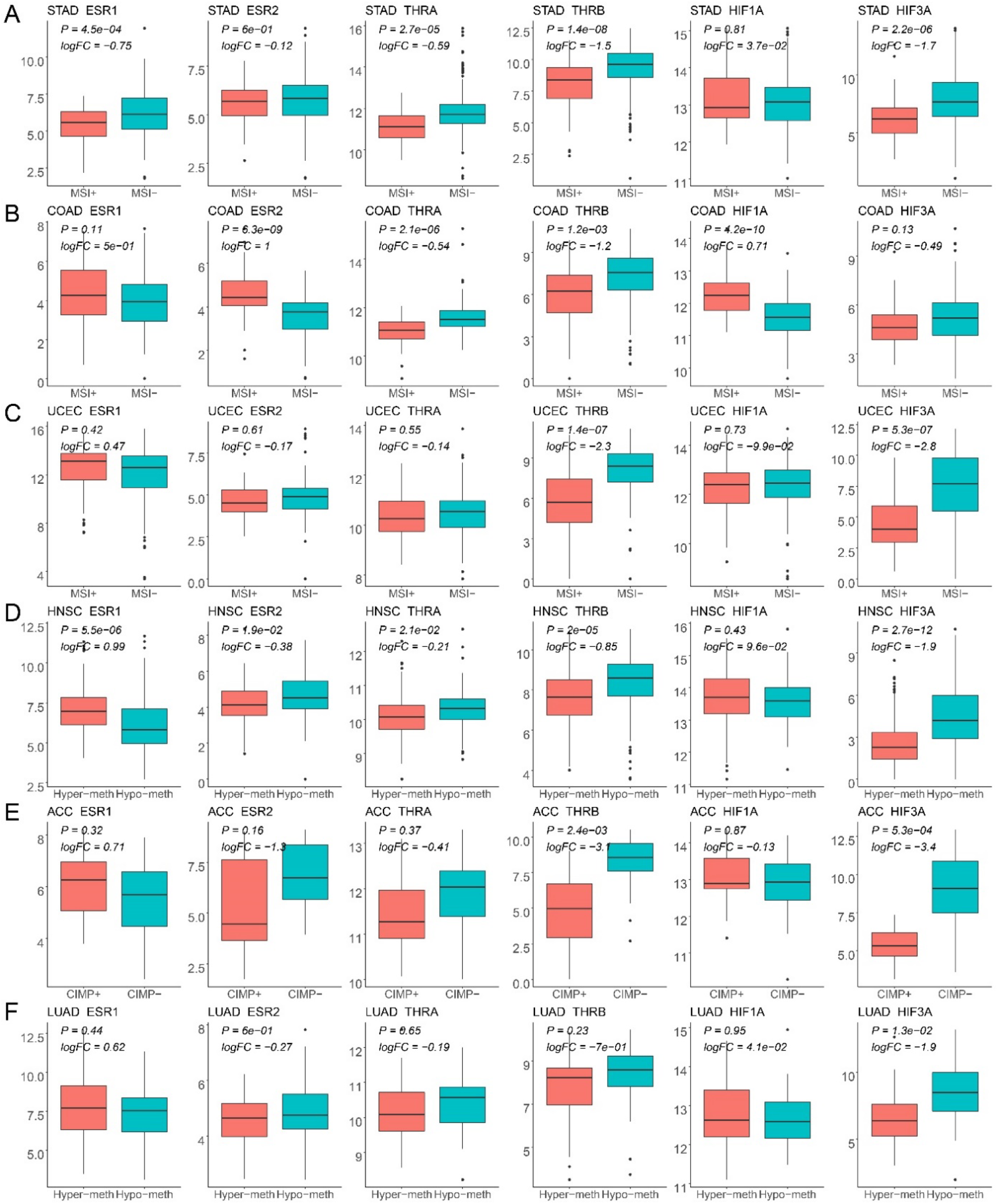
Boxplot matrixⅠof *ESR1*, *ESR2*, *THRA*, *THRB*, *HIF1A*, and *HIF3A*. Boxplots of the six genes mentioned in the main text, comparing gene expression between two groups: MSI+ versus MSI-, HyperMeth versus HypoMeth across STAD, COAD, UCEC, HNSC, ACC and LUAD. The adjusted P value and log fold change shown in the figure were obtained using Limma.

**Fig. S5.**
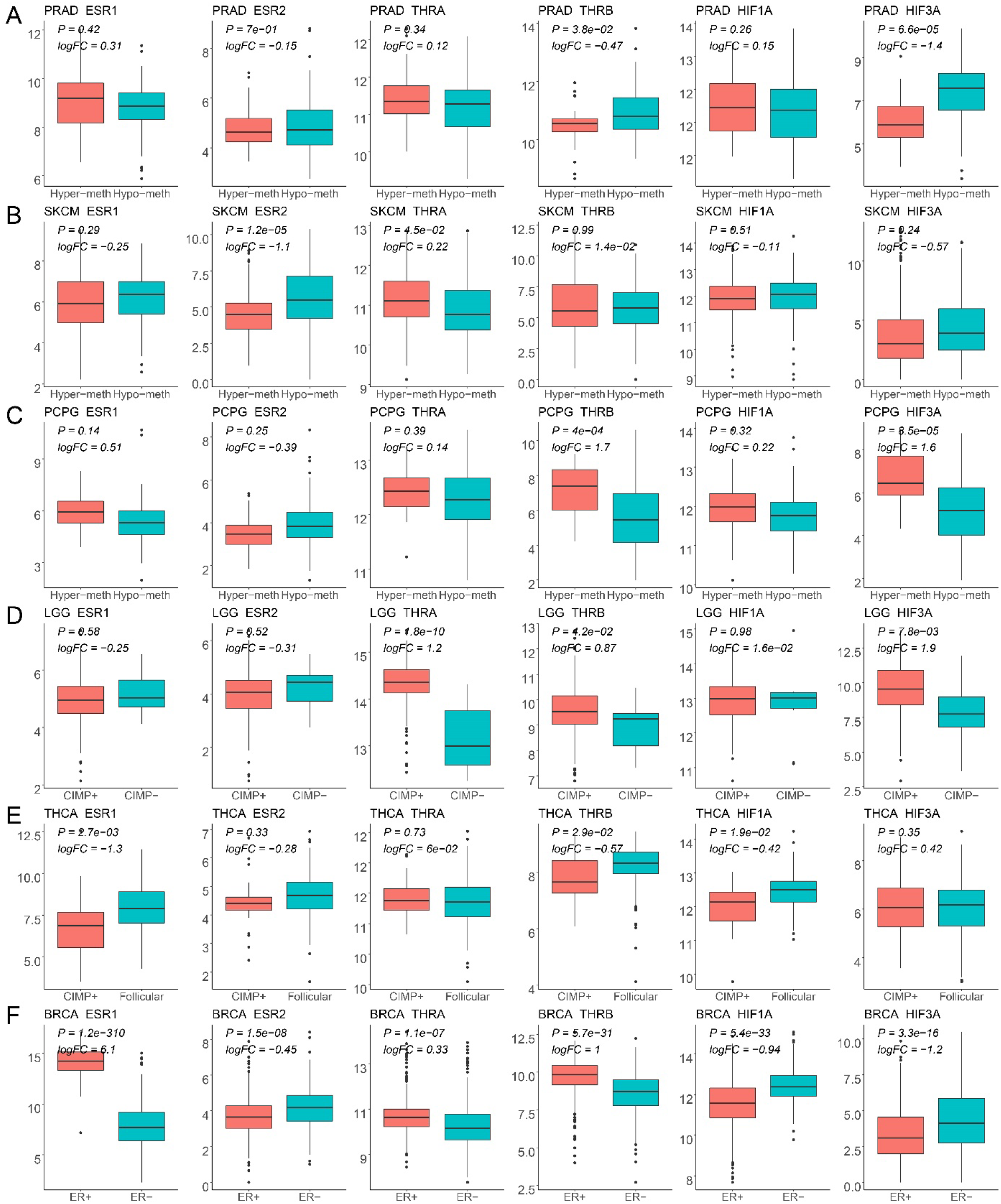
Boxplot matrixⅡof *ESR1*, *ESR2*, *THRA*, *THRB*, *HIF1A*, and *HIF3A*. Boxplots of the six genes mentioned in the main text, comparing gene expression between two groups: HyperMeth versus HypoMeth, and ER+ versus ER-across PRAD, SKCM, PCPG, LGG, THCA and BRCA. The adjusted P value and log fold change shown in the figure were obtained using Limma.

**Fig. S6.**
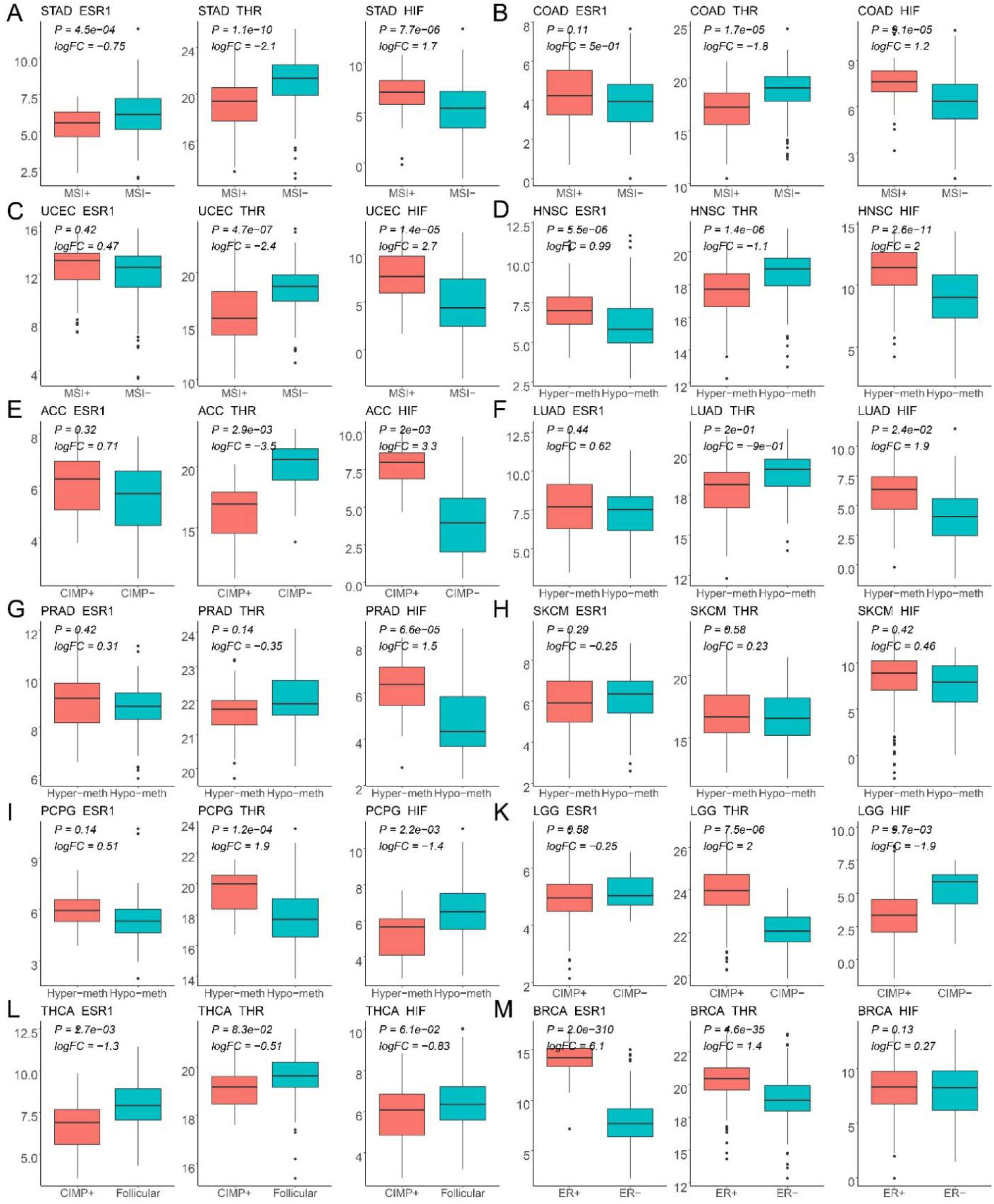
Boxplot matrix corresponding to Fig. 5. Each boxplot corresponds to each dot in Figure 5 and compares gene expression between two groups: MSI+ versus MSI-, HyperMeth versus HypoMeth, and ER+ versus ER-. The adjusted P value and log fold change shown in the figure were obtained using Limma. (A-M) Dot plots from STAD to BRCA.

**Fig. S7.**
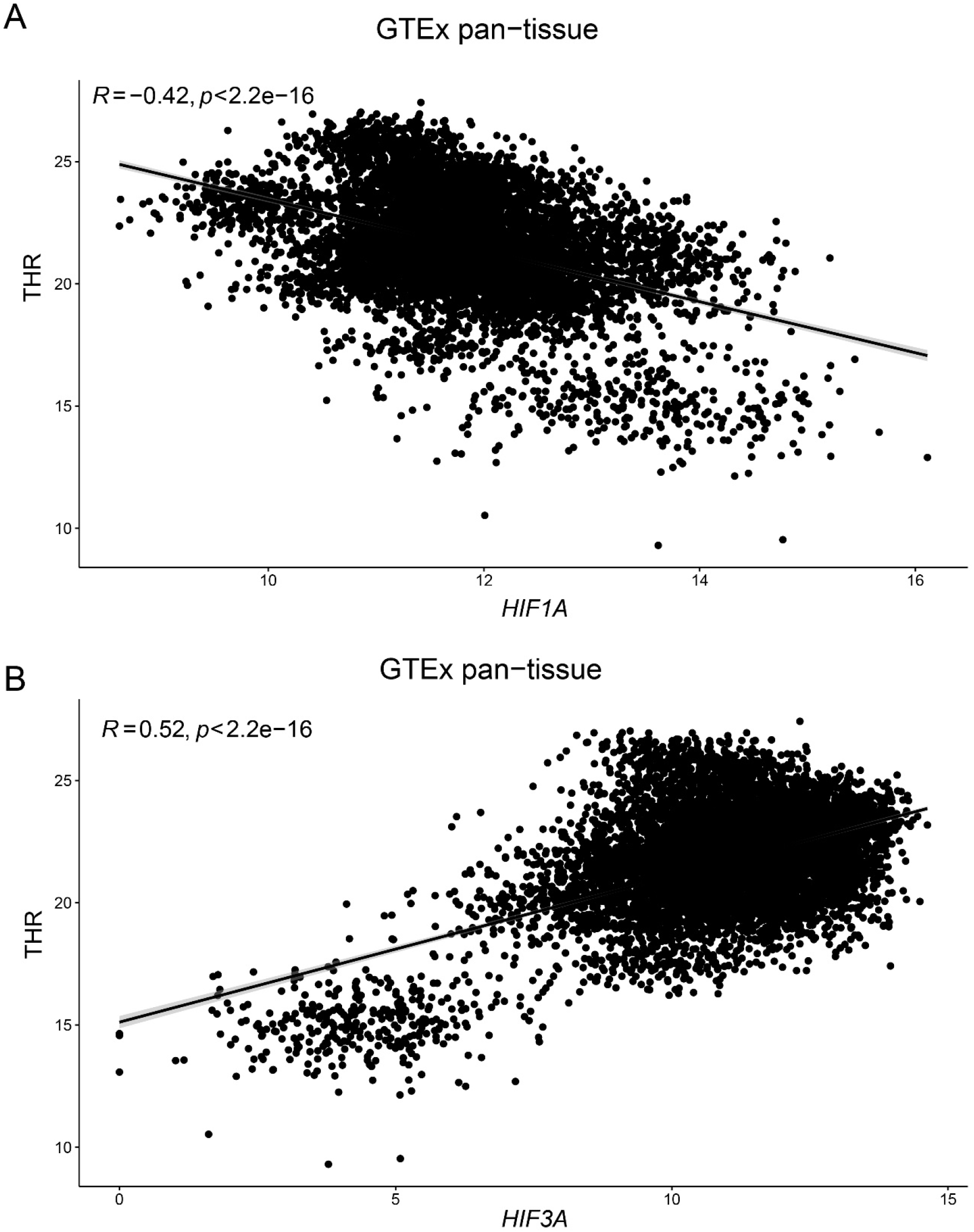
Correlations of THR with *HIF1A* and *HIF3A* in pan-tissue analyses. *R* represents the Pearson correlation coefficient; P values were obtained using Student’s t-test.

**Fig. S8.**
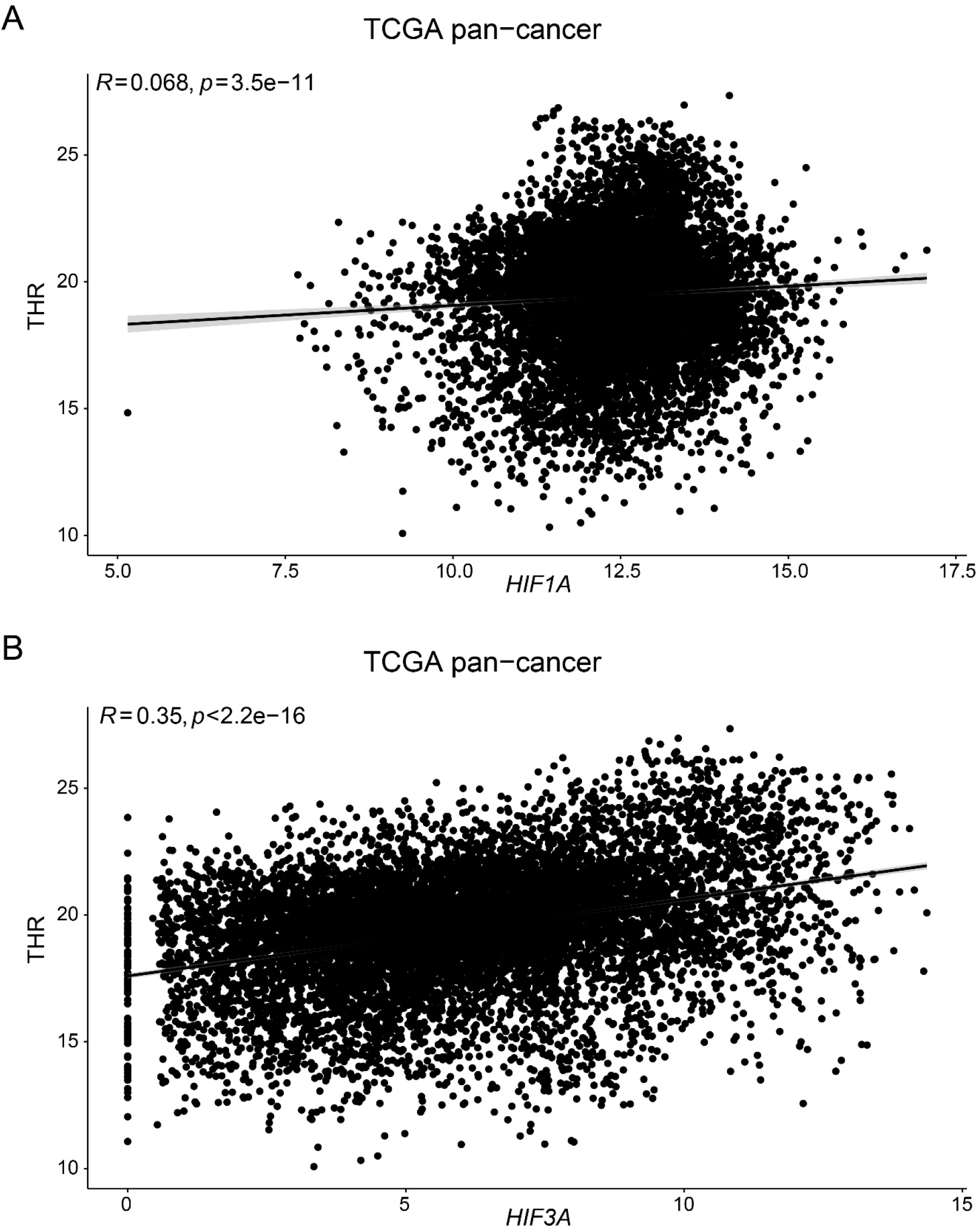
Correlations of THR with *HIF1A* and *HIF3A* in pan-cancer analyses. *R* represents the Pearson correlation coefficient; P values were obtained using Student’s t-test.

**Table S1.** Sample sizes for each tissue in GTEx data.

| **Tissue** | **Female** | **Male** | **Total** |
| --- | --- | --- | --- |
| Bladder | 3 | 6 | 9 |
| Kidney | 5 | 22 | 27 |
| Liver | 33 | 77 | 110 |
| Skin | 190 | 366 | 556 |
| Colon | 124 | 180 | 304 |
| Stomach | 73 | 99 | 172 |
| Pancreas | 72 | 93 | 165 |
| Lung | 100 | 187 | 287 |
| Thyroid | 105 | 173 | 278 |
| Brain | 353 | 783 | 1136 |
| Esophagus | 234 | 415 | 649 |
| Adrenal gland | 57 | 70 | 127 |

**Table S2.** Tissue-specific sex-biased expression of the GnRH secretion pathway based on GTEx data.

| **Tissue** | **P value** | **LogFC** |
| --- | --- | --- |
| Adrenal gland | 0.030899 | 0.100831 |
| Stomach | 0.012977 | 0.08919 |
| Thyroid | 0.009457 | 0.066973 |
| Pancreas | 0.112046 | 0.057273 |
| Skin | 0.068307 | 0.039208 |
| Colon | 0.414889 | 0.020896 |
| Lung | 0.565389 | 0.013314 |
| Esophagus | 0.533371 | 0.011119 |
| Kidney | 0.955948 | -0.00815 |
| Brain | 0.269388 | -0.02215 |
| Liver | 0.579539 | -0.03118 |

P values shown are adjusted P values obtained using Limma.

**Table S3.** Tissue-specific sex-biased expression of the GnRH signaling pathway based on GTEx data.

| **Tissue** | **P value** | **LogFC** |
| --- | --- | --- |
| Adrenal gland | 0.002109 | 0.100018 |
| Stomach | 0.221583 | 0.034591 |
| Thyroid | 0.167189 | 0.029276 |
| Pancreas | 0.409804 | 0.022836 |
| Brain | 0.885571 | 0.002091 |
| Colon | 0.683874 | -0.00639 |
| Esophagus | 0.196701 | -0.01705 |
| Lung | 0.402423 | -0.0175 |
| Skin | 0.217595 | -0.01834 |
| Liver | 0.482774 | -0.03411 |
| Kidney | 0.702808 | -0.03714 |

P values shown are adjusted P values obtained using Limma.

**Table S4.** Co-occurrence and mutual exclusivity analysis of the three hotspots.

| Hotspot1 | Hotspot2 | Co-occur<br>ring<br>samples | Hotspot1<br>-only<br>samples | Hotspot2<br>-only<br>samples | No<br>hotspot<br>samples | P value | Direction |
| --- | --- | --- | --- | --- | --- | --- | --- |
| *PTEN_267* | *PGM5_98* | 5 | 22 | 22 | 1162 | 2.2e-4 | Co-occur |
| *PTEN_267* | *LARP4B_163* | 5 | 22 | 20 | 1164 | 1.5e-4 | Co-occur |
| *PGM5_98* | *LARP4B_163* | 5 | 22 | 20 | 1164 | 1.5e-4 | Co-occur |

Fisher's exact test was performed on three pairs of the three hotspot mutations. All pairs tended to co-occur as indicated in the "Direction" column.

**Table S5.** Proportions of subtypes in TCGA data.

| Cancer | Sex | MSI+/<br>HyperMeth/<br>ER+<br>samples | Total<br>samples | Ratio | Relative<br>risk (f/m) | P value |
| --- | --- | --- | --- | --- | --- | --- |
| STAD | Male | 40 | 252 | 0.15873 | 1.923664 | 0.0013 |
|  | Female | 40 | 131 | 0.305344 |  |  |
| COAD | Male | 30 | 172 | 0.174419 | 1.235928 | 0.412533 |
|  | Female | 36 | 167 | 0.215569 |  |  |
| UCEC | Female | 124 | 507 | 0.244576 | NA | NA |
| LGG | Male | 132 | 135 | 0.977778 | 0.93905 | 0.063995 |
|  | Female | 101 | 110 | 0.918182 |  |  |
| GBM | Male | 10 | 17 | 0.588235 | 0.772727 | 0.760509 |
|  | Female | 5 | 11 | 0.454545 |  |  |
| ACC | Male | 7 | 16 | 0.4375 | 0.825397 | 0.830716 |
|  | Female | 13 | 36 | 0.361111 |  |  |
| THCA | Male | 13 | 42 | 0.309524 | 0.590237 | 0.145401 |
|  | Female | 19 | 104 | 0.182692 |  |  |
| SKCM | Male | 96 | 204 | 0.470588 | 0.962891 | 0.843433 |
|  | Female | 58 | 128 | 0.453125 |  |  |
| PCPG | Male | 46 | 60 | 0.766667 | 1.032609 | 0.893141 |
|  | Female | 57 | 72 | 0.791667 |  |  |
| HNSC | Male | 95 | 203 | 0.46798 | 0.702909 | 0.050884 |
|  | Female | 25 | 76 | 0.328947 |  |  |
| LUAD | Male | 35 | 56 | 0.625 | 1.134545 | 0.460331 |
|  | Female | 39 | 55 | 0.709091 |  |  |
| PRAD | Male | 182 | 333 | 0.546547 | NA | NA |
| BRCA | Female | 770 | 1043 | 0.738255 | NA | NA |

P values were obtained using z-test for proportions.

**Table S6.** Proportions of MSI subtypes in the work by Akagi et al (*24*).

| Cancer | Sex | MSI+<br>samples | Total<br>samples | Ratio | Relative<br>risk<br>(f/m) | P value |
| --- | --- | --- | --- | --- | --- | --- |
| Gastric | Male | 60 | 1250 | 4.8% | 1.63 | 0.013 |
|  | Female | 45 | 574 | 7.8% |  |  |
| Colorectal | Male | 186 | 5676 | 3.3% | 1.37 | 0.0026 |
|  | Female | 185 | 4138 | 4.5% |  |  |
| Endometrial | Female | 234 | 1389 | 16.8% | NA | NA |

For gastric and colorectal cancers, the proportions of MSI subtypes were significantly higher in women than in men. P values were obtained using z-test for proportions.

**Table S7.**
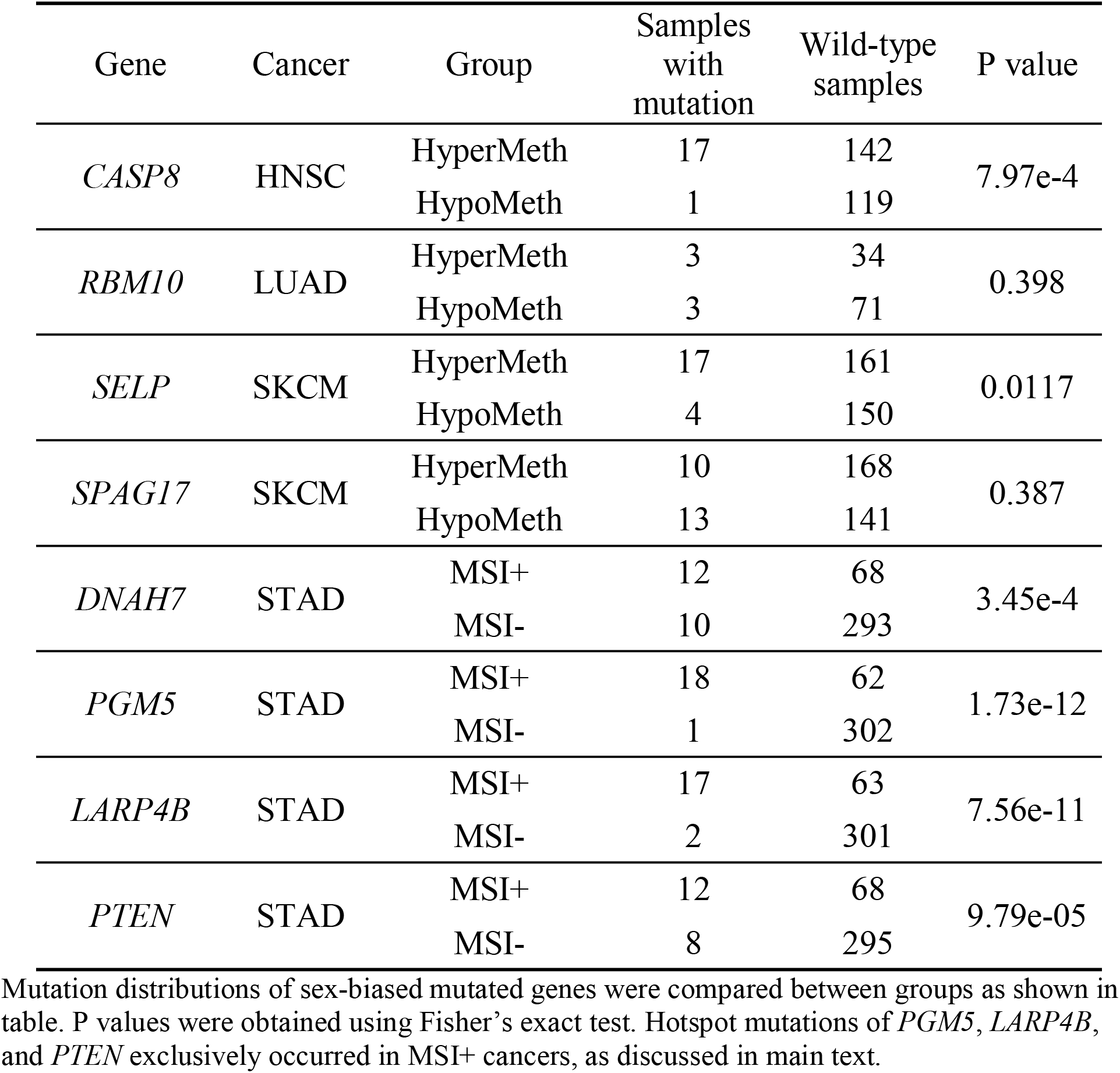
Mutation distributions of sex-biased mutated genes.

**Data S1.** Results of nonparallel pattern tests

**Data S2.** Results of sex-biased expressed pathway analysis in GTEx data

**Data S3.** Results of sex-biased mutated gene analysis in TCGA data

**Data S4.** Results of hotspot-grouped DEG analysis in STAD, COAD, and UCEC

**Data S5.** Results of subtype-grouped DEG analysis in TCGA data

